# "Same Difference": Comprehensive evaluation of three DNA methylation measurement platforms

**DOI:** 10.1101/077941

**Authors:** Thadeous J Kacmarczyk, Mame P. Fall, Xihui Zhang, Yuan Xin, Yushan Li, Alicia Alonso, Doron Betel

**Author notes:** Corresponding author: Thadeous J Kacmarczyk.

## Abstract

**Background:** DNA methylation in CpG context is fundamental to the epigenetic regulation of gene expression in high eukaryotes. Disorganization of methylation status is implicated in many diseases, cellular differentiation, imprinting, and other biological processes. Techniques that enrich for biologically relevant genomic regions with high CpG content are desired, since, depending on the size of an organism’s methylome, the depth of sequencing required to cover all CpGs can be prohibitively expensive. Currently, restriction enzyme based reduced representation bisulfite sequencing and its modified protocols are widely used to study methylation differences. Recently, Agilent Technologies and Roche NimbleGen have ventured to both reduce sequencing costs and capture CpGs of known biological relevance by marketing in-solution custom-capture hybridization platforms. We aimed to evaluate the similarities and differences of these three methods considering each targets approximately 10-13% of the human methylome.

**Results:** Overall, the regions covered per platform were as expected: targeted capture based methods covered >95% of their designed regions whereas the restriction enzyme-based method covered >70% of the expected fragments. While the total number of CpG loci shared by all methods was low, ~30% of any platform, the methylation levels of CpGs common across platforms were concordant. Annotation of CpG loci with genomic features revealed roughly the same proportions of feature annotations across the three platforms. Targeted capture methods encompass similar amounts of annotations with the restriction enzyme based method covering fewer promoters (~9%) and shores (~8%) and more unannotated loci (7-14%).

**Conclusions:** Although all methods are largely consistent in terms of covered CpG loci and cover similar proportions of annotated CpG loci, the restriction based enrichment results in more unannotated regions and the commercially available capture methods result in less off-target regions. Quality of DNA is very important for restriction based enrichment and starting material can be low. Conversely, quality of the starting material is less important for capture methods, and at least twice the amount of starting material is required. Pricing is marginally less for restriction based enrichment, and number of samples to be prepared is not restricted to the number of samples a kit supports. The one advantage of capture libraries is the ability to custom design areas of interest. The choice of the technique should be decided by the number of samples, the quality and quantity of DNA available and the biological areas of interest since comparable data are obtained from all platforms.

## Background

DNA cytosine methylation in the form of 5-methylcytosine (5mC) in CpG context is an epigenetic marker that is important for regulation of gene expression. Changes in CpG methylation are implicated in many diseases, and proper methylation patterns are required for normal development [1]. Large scale studies such as ENCODE [2] and the Human Epigenomics Roadmap [3] have performed extensive profiling of 5mC in various cell lines and tissues revealing a rich and dynamic landscape of 5mC patterns in the human genome. Given the importance of these markers to cellular development and contribution to disease, a number of approaches have been developed for detecting the methylation status of cytosines [4], with bisulfite sequencing (BS-seq) being widely used to provide single-base quantitative measurement of 5mC (and 5-hydroxymethylcytosine, 5hmC). Bisulfite sequencing refers to massively parallel sequencing after chemical deamination of cytosines (C) to uracils (U), followed by polymerase chain reaction (PCR). The deamination of cytosines is accomplished by the use of sodium bisulfite, and this pre-treatment preserves both the methyl modification in 5mC and the 5-hydroxymethyl modification in 5hmC [5]. The benchmark in methylome coverage is whole genome bisulfite sequencing (WGBS), which at 30X sequencing coverage, accounts for ~94% of all cytosines in the genome with 99.8% of them being CpG loci [6]. However, different WGBS library preparation protocols can bias region coverage. Since no method completely covers the methylome, and biologically relevant CpGs have been identified in known genomic features [1,7], developing focused assays considering these features is in demand with the caveat that these approaches will leave gaps in the methylome potentially excluding important CpGs.

There are several methods for acquiring DNA methylation levels and we investigated the characteristics of three currently widely used platforms: i) enrichment by enzymatic digestion (MspI) enhanced reduced representation bisulfite sequencing (ERRBS)[8], ii) capture based Agilent SureSelect Methyl-Seq (SSMethylSeq), and iii) capture based Roche NimbleGen SeqCap Epi CpGiant (CpGiant).

In this paper we present an analysis of the methylation pattern of the human lung fibroblast IMR-90 cell line profiled by each of the platform protocols, using two technical replicate libraries for ERRBS, and two libraries each for SSMethylSeq and CpGiant, one at the manufacturer’s suggested concentration and one at a reduced concentration (Table 1 and Additional file 1: Table S1). The capture libraries differ only in concentration of input material and are treated effectively as technical replicates. All libraries were sequenced to equivalent depth and compared to a library made using WGBS.

## Materials and Methods

### Cell growth and DNA preparation

IMR90 cells (American Type Culture Collection, Manassas, VA cat # CCL-186) were provided by Dan Hasson (Mount Sinai School of Medicine, New York, NY). DNA from 5x10^7^ cells was purified using the Gentra Puregene DNA kit according to manufacturer protocol (cat # 158389, Qiagen Valencia, CA). DNA was resuspended in TE, quantified using fluorometric quantification (Qubit 2.0 ThermoFisher Scientific Waltham, MA) and quality was assessed by running on a 1% agarose gel.

### ERRBS (digestion-based enhanced reduced representation bisulfite sequencing)

Two ERRBS libraries were prepared as described in Garrett-Bakelman, et al [8]. Briefly, 75 ng of DNA was digested with the methylation insensitive MspI enzyme (C^CGG). After end-repair, A-tailing, and adapter ligation with Illumina TruSeq adapters, the region corresponding to 84-334bp was size-selected as two fractions. Each fraction was subjected to overnight bisulfite conversion (55 cycles of 95°C for 30 sec, 50°C for 15 min) using EZ DNA methylation kit (Cat # D5002, Zymo Research, Irvine CA). Purified bisulfite converted DNA was PCR amplified using TruSeq primers (Illumina Inc. San Diego, CA) for 18 cycles of denaturing, annealing and extension/elongation steps using Roche FastStart (cat # 03 553 361 001) – 94°C for 20 secs, 65°C for 30 secs, 72°C for 1 min, followed by 72°C for 3 min. The resulting libraries were normalized to 2nM and pooled at the same molar ratio. Samples were clustered at 6.5pM on a V3 paired-end read flow cell and sequenced for 100 cycles on an Illumina HiSeq 2500.

### Agilent SureSelect Methyl-Seq (SSMethylSeq)

Two libraries were made according to the company’s specifications using 3ug and 1ug of DNA. DNA was sonicated using a Covaris S220 sonicator (Covaris, Woburn, MA) to obtain products of 150-200bp. DNA was then end-repaired, A-tailed and ligated with methylated adapters to create pre-capture DNA libraries. DNA Libraries were then hybridized to the RNA SureSelect Human methyl-seq capture library at 65°C for 16 hrs. Hybridized products were purified by capture with Strepdavin beads and then subjected to bisulfite conversion (64°C for 2.5hr) using the Zymo EZ DNA Gold kit (Cat # D5005, Zymo Research, Irvine CA). The bisulfite treated libraries were PCR amplified for 8 cycles with Agilent Taq to get the required amount of DNA library and then indexed by another 6 cycles of PCR amplification to create the final libraries. Libraries were clustered at 11pM on a V3 paired-end read flow cell and sequenced for 100 cycles on an Illumina HiSeq 2500.

### Roche NimbleGen SeqCap Epi CpGiant (CpGiant)

Two libraries were made using 1ug and 0.25ug of starting material, according to the company’s specifications. DNA was sonicated using a Covaris S220 sonicator (Covaris, Woburn, MA) to obtain products of 180-220bp. DNA was then end-repaired, A-tailed and ligated with methylated indexed-adapters to create pre-capture DNA libraries. The pre-capture libraries were bisulfite converted at 54°C for 1 hour using the Zymo EZ DNA Lightning kit (Cat # D5030, Zymo Research, Irvine CA). The bisulfite treated pre-capture libraries were PCR amplified with HiFi HotSart Uracil+ polymerase (Cat# KK2802, Kapa Biosystems, Wilmington, MA). The amplified, bisulfite converted sample libraries were then hybridized to the probe pool of fully-, partially-and un-methylated cytosines from both strands of DNA oligos at 42°C for 72 hrs. Hybridized products were purified by capture with Capture Beads and PCR amplified for 15 cycles to create the final libraries. Libraries were clustered at 12pM on a V3 paired-end read flow cell and sequenced for 100 cycles on an Illumina HiSeq 2500.

### Whole Genome Bisulfite Sequencing (WGBS)

Briefly, 100 ng of DNA were bisulfite converted using EZ DNA Methylation-Gold Kit (cat # D5005, Zymo Research Corporation, Irvine, CA) and the single stranded DNA obtained processed for library construction using the EpiGnome Methyl-Seq kit (Cat. # EGMK81324) as per manufacturer protocol (Illumina Madison, Madison, WI). The DNA was made double stranded by the use of 5’tagged random hexamers and subsequently 3’ tagged with terminal tagging oligo. The di-tagged DNA was enriched using 10 cycles of PCR, with PCR indexed-primers compatible with Illumina sequencing. The library was clustered at 7 pM on a V3 paired-end read flow cell and sequenced for 100 cycles on an Illumina HiSeq 2500.

### Computational analysis

Illumina’s CASAVA 1.8.2 was used to generate fastq files from basecalls. Raw fastq reads were processed by a custom pipeline that consists of: 1) filtering raw fastq reads for pass filter reads, 2) trimming adapter sequence by FLEXBAR [9], 3) genomic alignments performed using Bismark [10] and Bowtie [11] to reference human genome hg19, and 4) methylation calling by a custom PERL script [8]. Custom analysis scripts were written in R (version 3.3.0 [12]), including packages: Bioconductor - Biobase version 2.32.0 [13], GenomicRanges package version 1.24.0 [14], Beeswarm package version 0.2.3 [15], Affy package version 1.50.0 [16], BSgenome.Hsapiens.UCSC.hg19 package version 1.4.0 [17], PreprocessCore package version 1.34.0 [18], UpSetR package version 1.2.0 [19], and VennDiagram package version 1.6.17 [20], were used to perform the analysis and are available at GitHub (https://github.com/thk2008/methylseqplatformcomparison). Sequencing data and methylation results are available to download from GEO (GSE83595)

## Results

### Region coverage

Targeted capture techniques (SSMethylSeq and CpGiant) have a designed set of genomic regions and therefore, a predicted set of CpGs covered. SSMethylSeq is specifically designed to capture CpGs from a single DNA strand, where the other platform capture CpGs from both DNA strands. The ERRBS protocol is considered targeted to the extent that the restriction digest produces consistent genomic fragments, sizes from 84-334bp are isolated during the library preparation gel extraction step. Note that since the DNA for WGBS is randomly sheared and coverage depends upon sequencing depth, there are no predicted regions per se and consequently WGBS is excluded from the region coverage analysis. Regions are considered overlapping if any two regions overlap by at least 10 bases. Overall, SSMethylSeq and MspI 84-334b have similar region length proportions, while CpGiant has fewer regions and more variable lengths (Figure 1A and Additional file 1: Table S2). Additionally, the two capture platforms cover similar regions, and similar amount of different regions from MspI 84-334b (Figure 1B).

**Figure 1.**
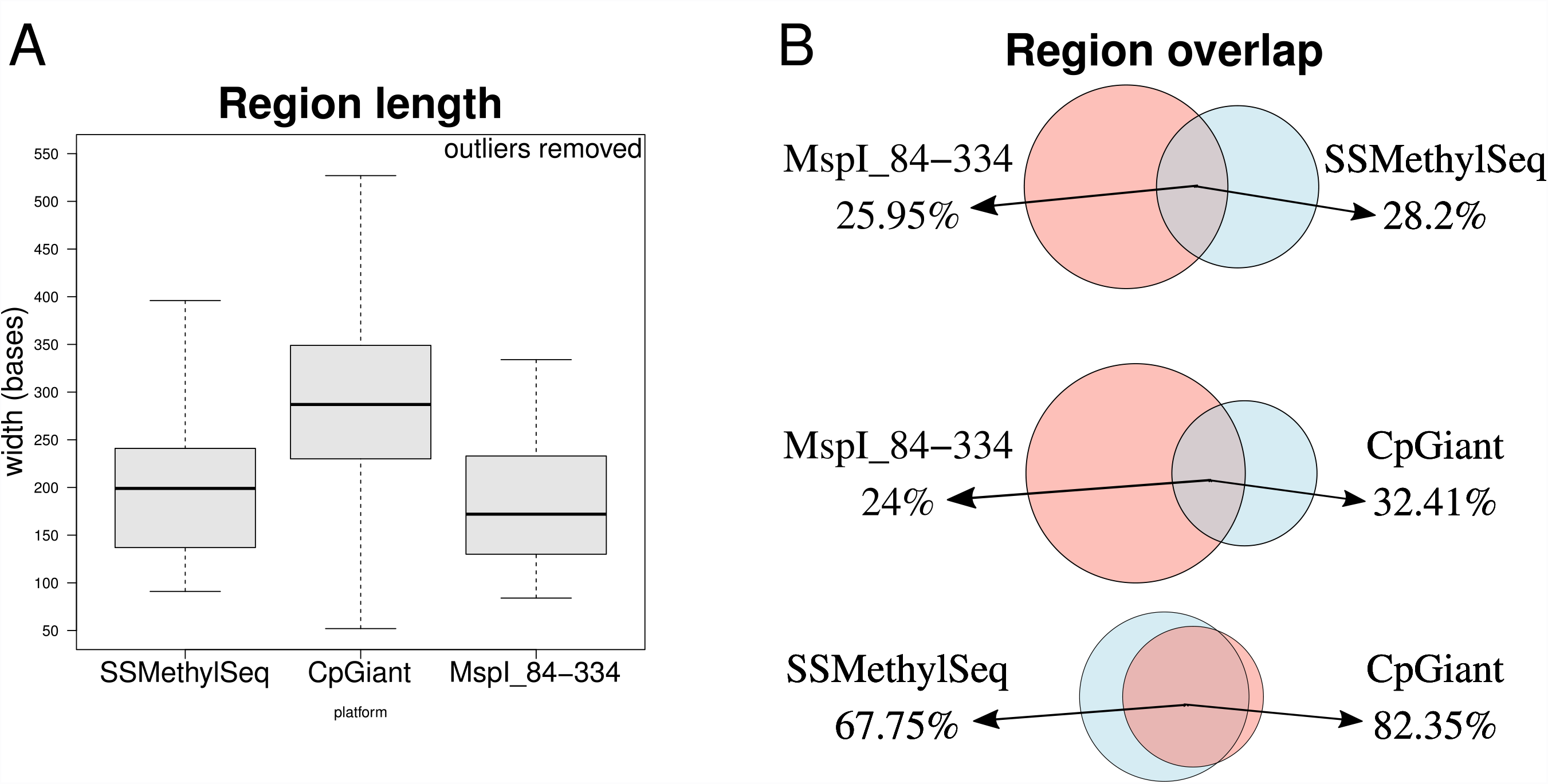
Length and overlap of design regions and MspI predicted regions. **A**) Boxplot showing the distributions of targeted region’s lengths. SSMethylSeq and MspI 84-334b (ERRBS) show similar region proportions, while CpGiant regions are generally longer and are more variable in length. **B**) Venn diagrams showing the pairwise overlap comparison of each of the platform’s regions and the total number of regions. Circle size is proportional to the size of the set. The two capture platforms have a higher degree of common regions with each other than either one with MspI 84-344b.

### Description of analysis

Even though the platforms are different, the data are all bisulfite sequencing data, thus the procedure for processing the data is the same. First we evaluated sequencing and alignment consistency. Next we compared strand symmetry of methylation values, since the SSMethylSeq platform is designed to cover predominantly one strand. Then we evaluated several measures to quantify properties of each platform relating to the target regions that are covered, methylation levels of cytosines in CpG context and coverage of genomic regions. Target region analysis includes the number of CpGs covered, the fraction of target regions covered, the coverage of target region CpGs, the overlap of CpGs across all platforms, and the concordance of methylation levels of overlapping CpGs. Similarly, genomic region evaluation is comprised of the number and fraction of annotated CpGs by genomic feature (i.e. CpG islands or shores – 2kb flanking the islands) and by gene part (i.e. promoter, exon, intron), and the overlap, coverage, and concordance of genomic annotations. We used the median absolute deviation (MAD), a robust measure of variability insensitive to outliers, to estimate the statistical dispersion in methylation level comparisons.

### Sample preparation and sequencing

The general library preparation protocol for ERRBS is to digest input DNA with the methylation insensitive MspI enzyme followed by addition of adapters and barcodes, enrichment by gel size selection for fragments 84-334bp, bisulfite conversion, and amplification by PCR. The capture methods of SSMethylSeq and CpGiant are similar in that targeted capture of genomic regions is performed by hybridization to probes thereby providing a direct measure of CpG methylation at predefined regions. However, the methods differ in two important aspects. First, the SSMethylSeq platform captures and measures methylation from only one DNA strand. Second, in CpGiant’s methodology bisulfite conversion is performed before hybridization to oligonucleotides whereas in SSMethylSeq bisulfite conversion is performed prior to genomic capture. Hence, CpGiant’s probe design must capture the various combinations of DNA sequences that can result from bisulfite conversion. In contrast to the targeted approaches WGBS is unbiased and relies on bisulfite breakage of genomic DNA. All libraries were made from human lung fibroblast IMR90 cell line DNA and prepared as described in the materials and methods with prominent differences outlined in Table 1.

**Table 1:** Protocols comparison

|  | **ERRBS** | **SS-MethylSeq** | **CpGiant** | **WGBS** |
| --- | --- | --- | --- | --- |
| DNA requirement | 75ng >40kb | 1, 3ug | 0.25,1ug | 100 ng >40kb |
| DNA processing | MspI digestion to completion followed by fractionation of 84-334bp | Sonication to 150-200bp | Sonication to 180-220bp | Single stranded and fragmented during bisulfate conversion |
| Enrichment method | Size fractionation of 84-334bp sizes | Hybridization to RNA capture library stranded design | Hybridization to oligo probes containing fully-, partially- and un-methylated cytosines from both strands | none |
| Bisulfite conversion step | Post-adapter ligation | Post-hybridization capture | Pre-hybridization capture | Pre-adapter ligation |
| Zymo Research Bisulfite Conversion kit | EZ-DNA Methylation (50°C, 55cycles) | EZ-DNA Methylation Gold (64°C, 2.5hr) | EZ-Methylation Lightning (54°C, 1hr) | EZ DNA Methylation-Gold |
| Total PCR amplification cycles | 18 Post enrichment and bisulfite conversion | 16 Post enrichment and bisulfite conversion (8 for amplification and 6 for Indexing) | 28 12 post bisulfite conversion & 16 post enrichment | 10 Post bisulfite conversion |
| DNA Polymerase (uracil tolerant) | FastStart Taq (Roche) | Taq2000 (Agilent Technologies) | HiFi HotSart Uracil+ (Kapa Biosystems) | FailSafe Enzyme (Epicentre) |
| Predicted number of targeted CpG | 6.6M | 3.7M | 5.6M | 56M |

| sites |  |  |  |  |
| --- | --- | --- | --- | --- |
| Relative price<br>(library<br>preparation and<br>fixed coverage) | 13.5 % | 15.5% | 15.5% | 100% |

### Sequencing and alignment characteristics

Generally, all platforms produced similar sequencing results with no noticeable bias or reduced quality scores. The number of clusters and number of pass filter reads produced a typically consistent number of usable reads for all samples (254M +/− 37M), see Additional file 1: Table S3 for more details.

Consistent with other bisulfite converted samples the number of uniquely mapped reads was ~ 70.8% sd=5.2% (Figure 2A). Across all platforms, a mean of 24.2%, s= 2.2% of reads were not aligned and ambiguously mapped reads showed a larger proportion for ERRBS (~12.1%), than WGBS (4.6%), SSMethylSeq (1.4%), or CpGiant (1.4%) (Figure 2A). These observations indicate the propensity of designed platforms to avoid repetitive regions or paralogous regions in contrast to ERRBS where no such selection is possible.

**Figure 2.**
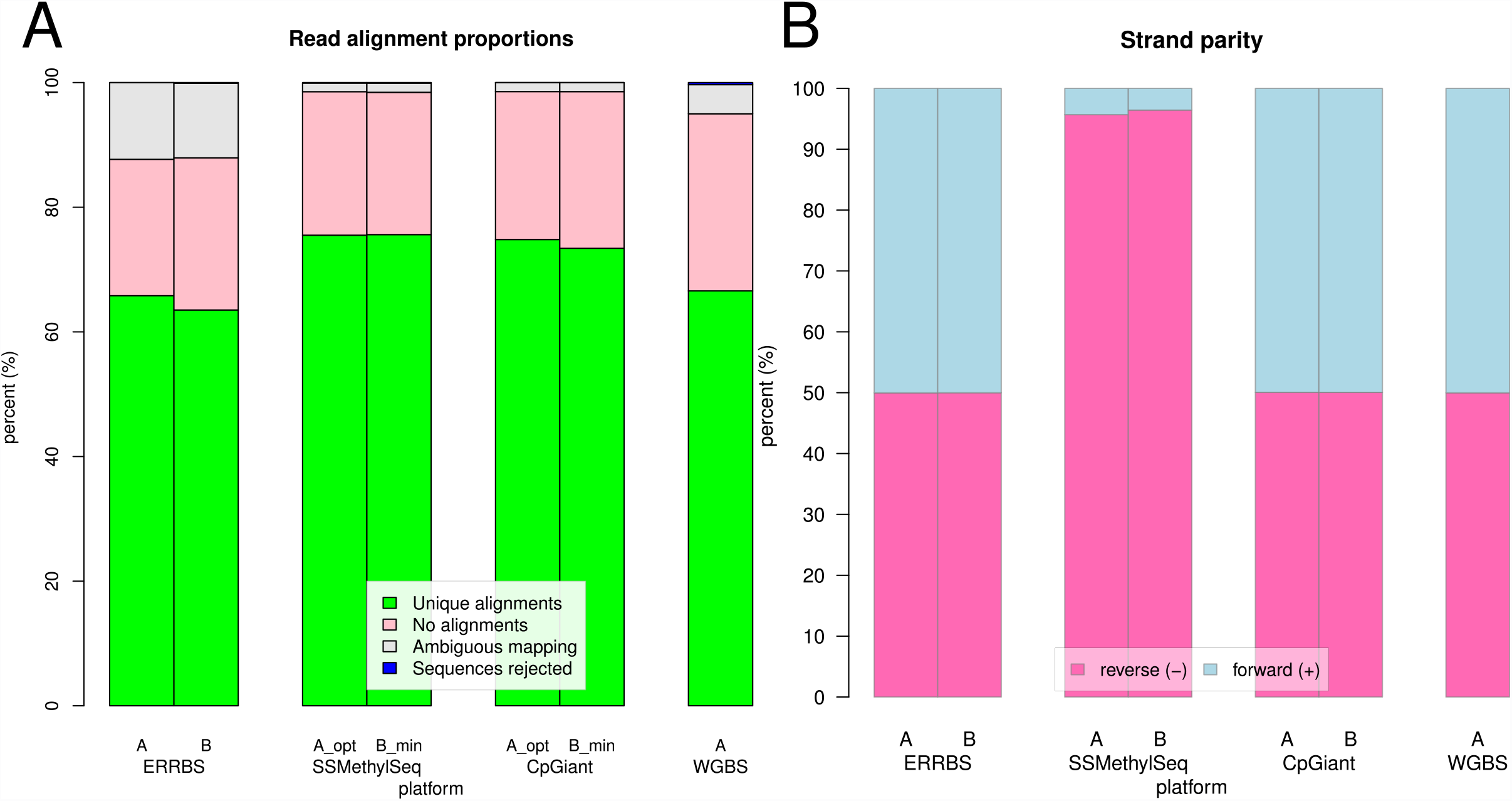
Read alignment and strand parity. **A**) Percent alignment of uniquely aligned reads (green), ambiguously mapped reads (grays), reads with no alignment (pink) and rejected reads (blue). **B**) The fraction of CpG’s covered at >=10x coverage grouped by strand, plus strand (blue) and the minus strand (pink). The strand specific protocol of SSMethylSeq platform is evident by the high proportion of reads mapped to the negative strand.

### Strand methylation symmetry

Maintenance of symmetric methylation patterns across complementary CpG sites is required in order to preserve methylation patterns across cellular divisions. In principle, measuring methylation levels on one strand is sufficient to infer the cellular methylation state. Indeed, the SSMethylSeq platform is designed to capture only one strand in contrast to all other platforms that capture from both strands (Figure 2B). We note that this is different from measuring hemi-methylation states, which refers to different methylation between two parental alleles. This requires genomic phasing information and is not considered in this analysis. To validate that methylation is indeed strand symmetric we compared the methylation values between complementary CpG sites (Figure 3). For ERRBS and CpGiant about 40% of the CpGs were complementary and analysis of SSMethylSeq was done with the ~2% of the probes that were complementary. We found strong agreement in methylation values in all samples and all platforms supporting the notion of symmetric methylation (Figure 3, mean MAD=0.28, sd=0.06). ERRBS contains a small set of discordant CpG sites (Figure 3A, B), where methylation values are inconsistent among complementary sites (Δ > 99, 3.8% of sites ERRBS_A, 1.0% in ERRBS_B at 10X coverage). This discordance is a consequence of the ERRBS library preparation where the MspI staggered cut sites are *de novo* (*in vitro*) filled with C’ and G’ to generate blunt ends. As a result, methylated CpGs at the restriction sites (i.e. at the ends of sequences fragments) are discordant. To correct this artificial discordance, it’s common for single end sequencing to discard CpG values at MspI cut sites. We observed additional disparity occurring in paired-end sequencing where the sequenced fragments are shorter than the overall sequencing length where sequencing reads from opposite strands are overlapping. Since the fraction of discordant sites is small, we have kept them in the analysis. However, In order to maintain equal evaluation of CpG capture and methylation levels among all platforms, all subsequent analyses are based on CpG-units, which are defined as CpG sites with >=10 spanning reads regardless of which strand the reads are mapped (i.e. if two CpG sites are complementary they are combined into one unit and then filtered for >=10x coverage).

**Figure 3:**
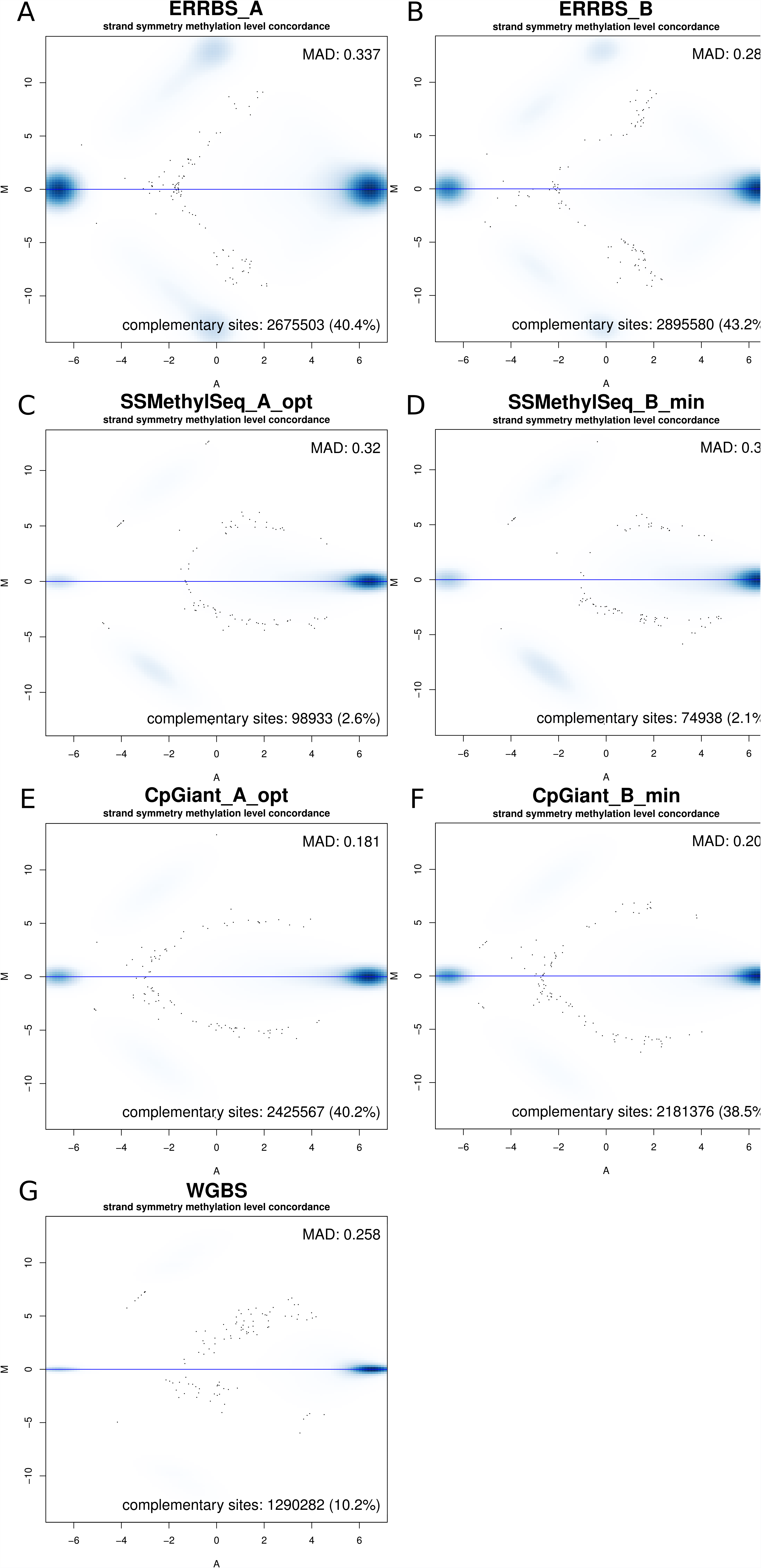
Strand symmetry of methylation values MA-plots. MA-plots of the log average of the methylation levels (A) on the x-axis and log ratio of the methylation levels (M) on the y-axis, between complementary CpG positions. Median absolute deviation (MAD) values are used to evaluate the agreement in methylation levels. The bimodal nature of methylation patterns (mostly unmethylated or methylated) is reflected in the high density at both ends of the x-axis. The artificially discordant sites introduced during ERRBS library preparation are identified as increased density off the center line at the low methylation values (at A < 0 range) in panels A, B. Panels are: **A**) ERRBS_A, **B**) ERRBS_B, **C**) SSMethylSeq_A, **D**) SSMethylSeq_B, **E**) CpGiant_A, **F**) CpGiant_B, and **G**) WGBS.

### CpG-unit coverage and target region coverage

We evaluated the number of CpG-units covered and the mean coverage of all CpG-units that were sequenced at 10X depth or more. ERRBS, SSMethylSeq and CpGiant platforms cover on average 3.8M CpG-units, sd=0.2M at mean coverage of 114.27, sd=23.6 (Figure 4A,B and Additional file 1: Figures S1 and S2). In contrast, WGBS sequenced at ~400M paired-end reads covered 14.4M CpG-units (~50% of the 28M total CpG-units in the genome) at a mean coverage of 19x, demonstrating that achieving comparable coverage by WGBS is significantly more costly than targeted platforms. Therefore, these results confirm that while WGBS profiles a large portion of CpG-units, the targeted platforms provide a cost effective way to interrogate a limited, yet potentially most informative, set of CpG-units at considerably reduced cost.

**Figure 4.**
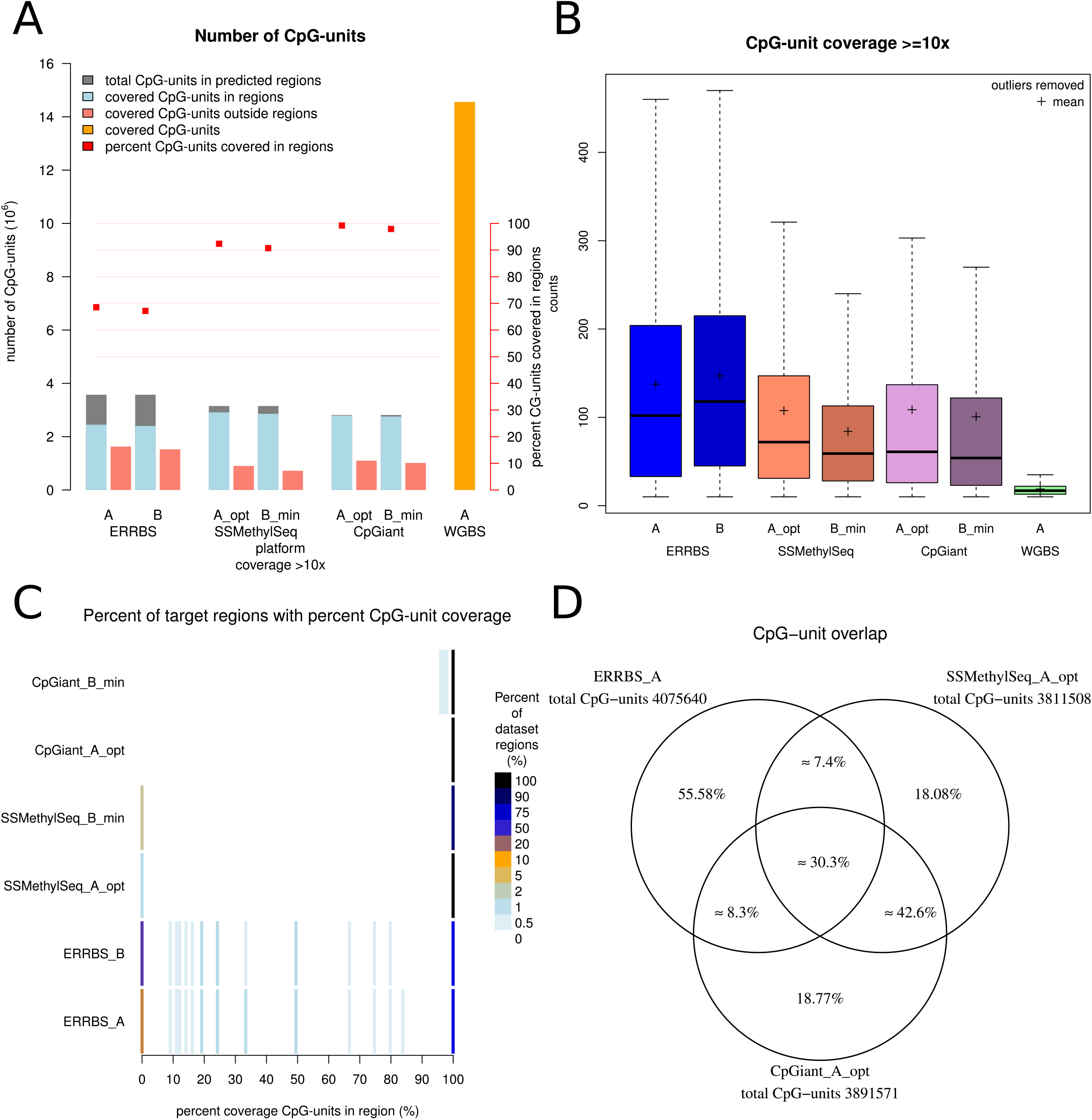
Platform CpG-unit region coverage and CpG-unit overlap. **A**) Number of CpG-units identified in targeted and off target regions by each platform. WGBS (orange) covers ~14.4M CpG-units however there is no notion of targeted regions in this platform. The targeted platforms predicted total CpG-units are depicted as gray bar and coverage of the CpG-units in the predicted regions (a.k.a “on-target”) are shown in blue. CpG-units outside the predicted set (off-target) are shown in red bars. The red square points and right scale represent the percent recovery of targeted CpG-units. **B**) Distribution of coverage per CpG-unit. **C**). Density plots of the percent of the dataset’s regions (color scale) with the fraction of CpG-units covered per region. These plots demonstrate that the reduced number of recovered CpG-units in ERRBS relative to the other platforms (red squares in panel A) are attributed to increased number of missed CpG-units shown as increased density at 0 region. **D**) Triple Venn diagram showing the proportion of overlap of CpG-units for the three platforms; size is not proportional to dataset.

Next, we evaluated the extent of coverage in the targeted regions and the fraction of their CpGs identified. Here, a region is considered as having coverage if at least one CpG-unit is sufficiently covered within the region. We observed that ERRBS covers on average 79.8% of its predicted regions 84-334bp, whereas CpGiant and SSMethylSeq cover, on average, 98.7% and 95.3% of their expected capture regions, respectively (Figure 4A). While region coverage and the coverage of CpG-units within the targeted regions are high, we observed that roughly 20%-40% of the CpG-units covered are outside the targeted regions (Figure 4A). In the case of ERRBS this is expected since restriction digestion and gel isolation can be variable. In the case of capture methods, this may indicate either cross-hybridization of the probes to other genomic locations or hybridization of longer fragments. Since there are typically several CpG-units within a region, we looked at the distribution of the fraction of CpG-units covered per region and found that nearly all CpG-units in targeted regions are covered although for both SSMethylSeq and ERRBS a number of CpGs in targeted regions are not represented presumably due to coverage bias (Figure 4C).

### CpG-unit overlap and methylation levels concordance

Overall, the average number of common CpG-units covered by all three platforms is ~30% +/− 1% (Figure 4D). When comparing the pairwise overlap of shared CpG-units across all samples, we observe high overlap between intra-platform replicates with median number of shared CpG-units ~3.58M, median overlap 92.1% and high methylation level concordance with mean MAD = 0.23 and sd = 0.05 (Additional file 1: Figure S3 and Table S4). Among inter-platform samples we observed less overlap; median number of shared CpG-units ~1.5M, median overlap 39.1% (Figure 5 upper triangle, Additional file 1: Figure S3 and Table S4). Methylation levels of common CpG-unit’s across all platforms are highly concordant indicating that methylation level measurements are consistent and reproducible with average MAD = 0.41 and sd = 0.25 (Figure 5 lower triangle, Additional file 1: Figure S3 and Table S4). Inter-platform concordance was slightly lower than intra-platform concordance with mean MAD = 0.46 and sd = 0.27 (Figure 5 lower triangle, Additional file 1: Figure S3 and Table S4). These results demonstrate that while the platforms differ in their capture approaches and bisulfite conversion kits (Table 1) these differences are not biasing methylation measurements. The differences among the platforms, therefore, are largely restricted to variations in targeted regions and not in methylation measurements.

**Figure 5.**
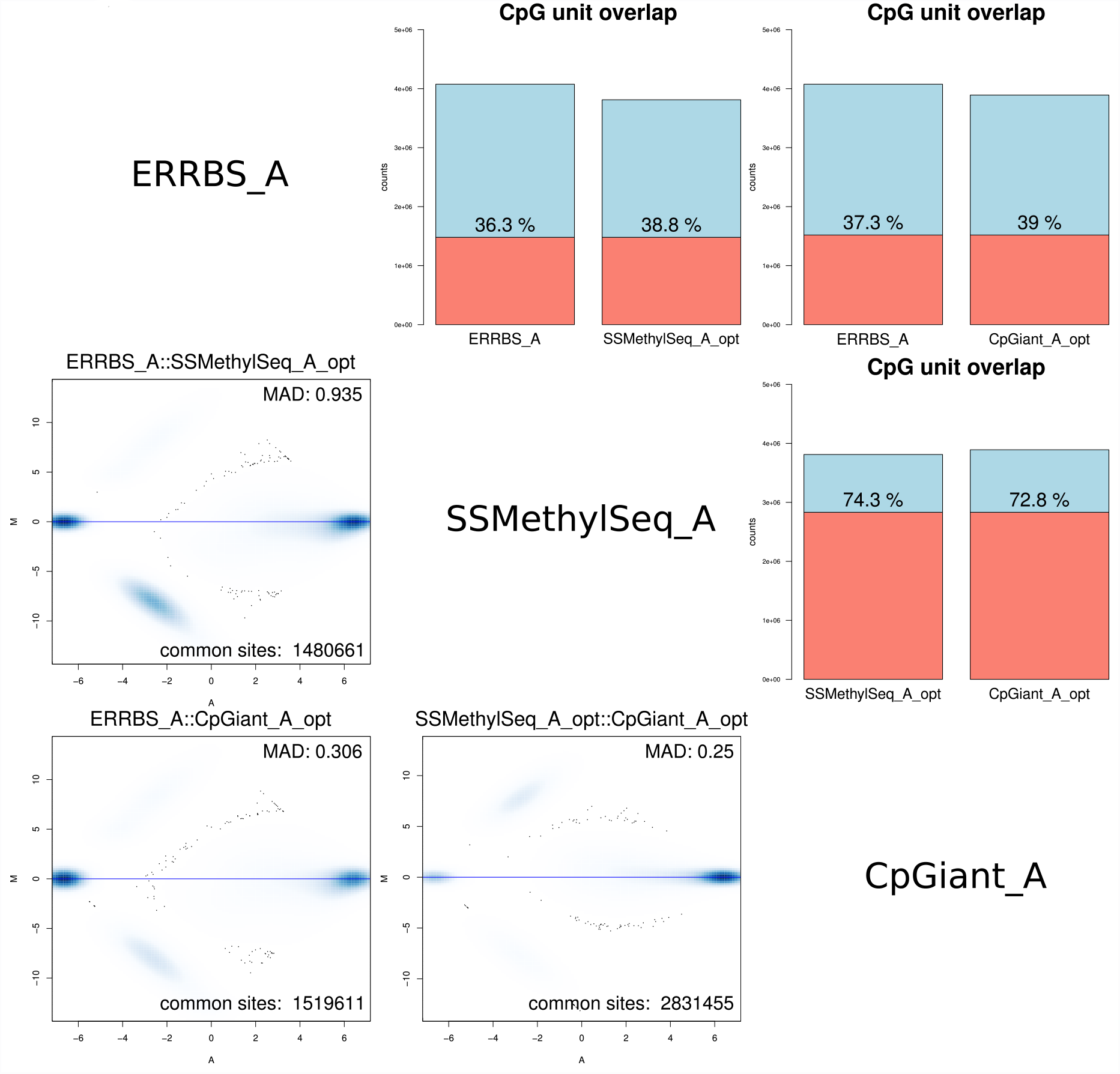
Inter-platform CpG-unit overlap and methylation levels concordance. The upper triangle shows the total number of CpG-units (blue) and number of overlapping CpG-units between any two samples (red) with the percent overlap indicated. The lower triangle shows MA-plots of common CpG-units between any two samples, where M is the log ratio of the methylation levels and A is the average log methylation level from the two platforms. There are blue clouds that at log scale show the variance in methylation at low levels. See Additional file 1: Figure S3 for all pairwise comparisons.

### Coverage of genomic feature regions by CpG-units

Next we were interested in what genomic features are covered by each platform and the degree of coverage. Analogous to the previous analysis, here we define a region as being one of exon, intron, promoter, CpGi, or shore, and coverage is a genomic region that contains at least one detected CpG-unit. It should be noted that the genomic feature annotations are not mutually exclusive and that some CpG-units are annotated by more than one category. Naturally, the targeted platforms focus on genomic regions known to play important roles in epigenetic regulation such as promoter regions, CpG islands, and CpG shores and ERRBS covers the same regions, but to a lesser degree (Figure 6A). Moreover, each platform covers similar proportions of CpG-units in particular genomic region (Figure 6B). Conversely, we looked at the number of CpG-units that have an annotation and observed similar trends for the three platforms with ERRBS having a larger proportion of unannotated CpG-units (~27%, Figure 6C). Overall, no platform is significantly enriched for particular genomic region although ERRBS is slightly less represented in promoters, CpG islands and CpG shores, while having more representation of unannotated CpG-units.

**Figure 6:**
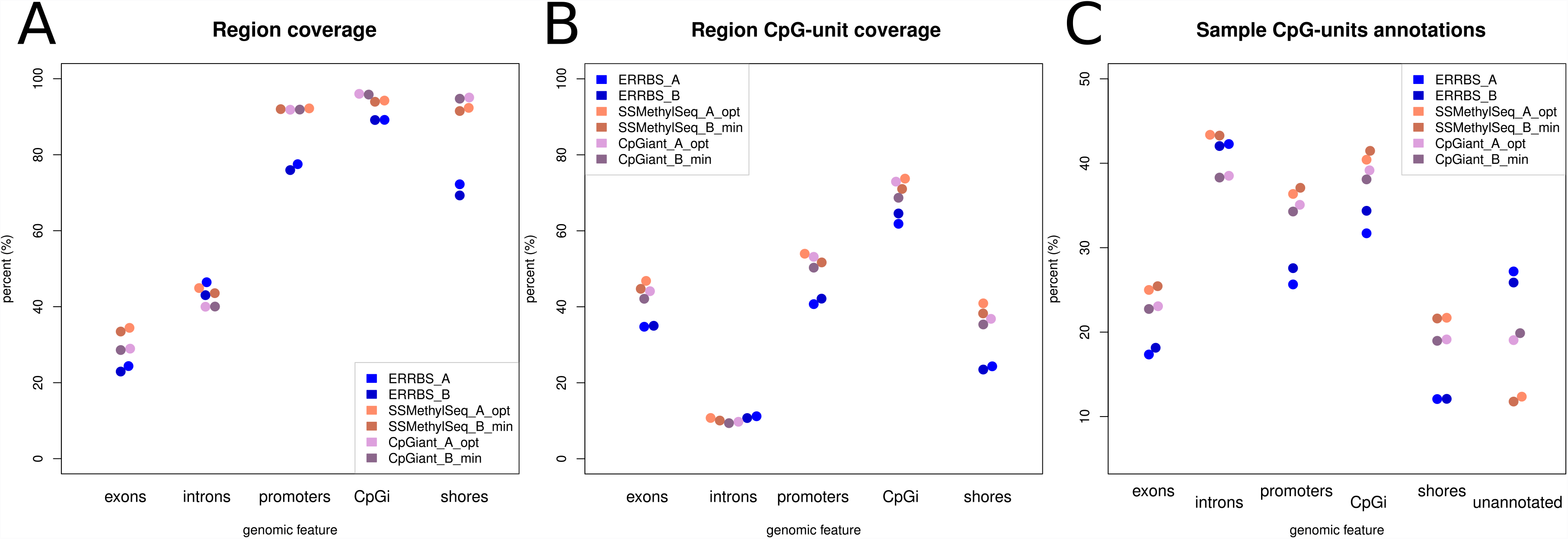
Summary of coverage and representation of annotated genomic regions by each platform. **A)** The fraction of genomic feature covered by sample CpG-units where at least one CpG-unit is covered in a region. **B)** The fraction of a region’s CpG-units covered, and **C)** The fraction of CpG-units annotated with a genomic feature. CpGi (CpG island).

### Overlap of CpG-units annotated with a genomic feature

We evaluated the overlap of annotated CpG-units to determine whether any platform is enriched for a particular genomic feature (e.g. exons, introns, promoters, CpG islands and shores). Again, it should be mentioned that the genomic feature annotations are not mutually exclusive and that some CpG-units are annotated by more than one category. However, a CpG-unit is counted as annotated if it has one or more annotations. Comparing the overlap in annotations across platforms, we see a similar grouping as above with the methylation levels, intra-platform overlap is high (mean overlap 93.7% sd=2.9%), and inter-platform overlap is lower (mean 55.1% sd=15.5%) (Figure 7 and Additional file 1: Figures S4-S9). Thus, we observed similar proportions of genomic region coverage across all platforms, but lower overlap of annotated CpG-units suggesting coverage of different loci within those regions.

**Figure 7:**
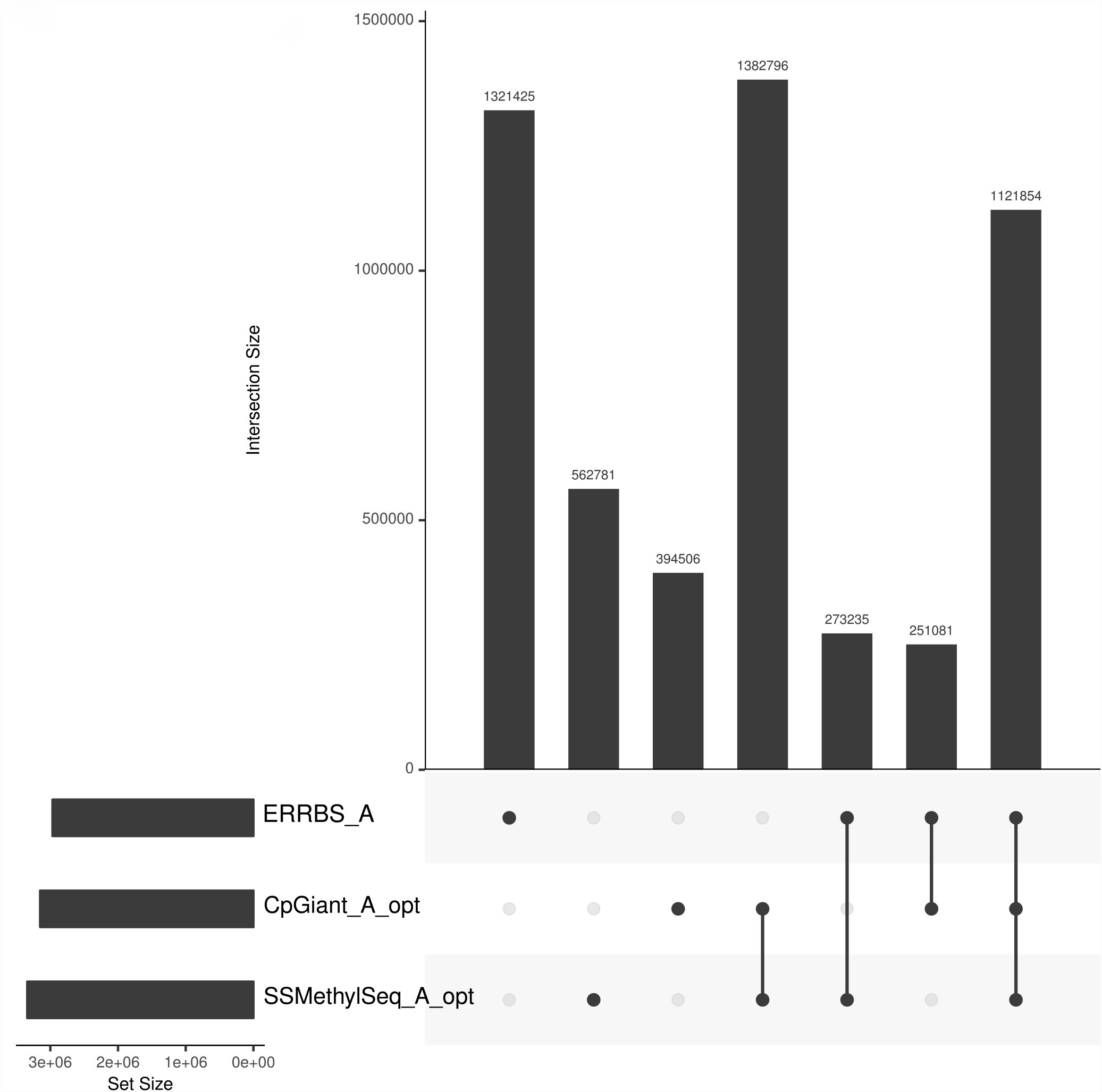
CpG-unit annotations overlap. Overlaps of all CpG-unit annotations across all platforms. The UpSet visualization technique [19] for set intersections is displayed as matrix layout. Horizontal bars on the lower left indicate the total number of annotated CpG-units in the set. Dark circles in the matrix (lower right) indicate sets that are part of the intersection. Bars in the main plot area (upper right) indicate the number of intersecting CpG-units for the sets represented by the dark circles. Roughly 33% of the annotations are common to all three suggesting that while there may be similar proportions of CpG-unit annotations (i.e. they may be covering similar regions), they are covering different loci within those regions.

## Discussion

We performed a systematic analysis of the characteristics of three prominent platforms for measuring DNA methylation levels: ERRBS, Agilent SureSelect Methyl-Seq (SSMethylSeq), Roche NimbleGen SeqCap Epi CpGiant (CpGiant), and WGBS with the goal of identifying whether one method outperforms the others for any particular property.

We assessed the expected region coverage for SSMethylSeq and CpGiant and MspI (for ERRBS) and observed that, overall, the CpG sites covered by each platform are largely distinct from each other. SSMethylSeq and CpGiant cover roughly 70% of the same CpGs, while MspI has only ~30% overlap with either capture. However, each platform covers a large fraction of its targeted regions. SSMethylSeq and CpGiant cover greater than 95% of their designed capture regions and ERRBS covers at least 77% of its expected fragments. Furthermore, methylation levels of overlapping CpG-units are highly concordant.

Finally, each platform covered roughly the same proportions of genomic features (CpG islands, shores, promoters, exons, and introns), but profiled different CpG sites within those regions. In addition to the differences in the targeted CpG positions, there are also differences in the library preparation protocols. ERRBS can be performed using small amounts of starting material (as little as 75ngs), whereas SSMethylSeq requires 1ug and CpGiant 0.25ug of starting DNA (although those are expected to reduce with further optimization by the vendors). ERRBS, or digestion-based methods, can cover certain genomic regions that are not amenable to capture. ERRBS provides comparable data to the other platforms, although there is more variability among the profiled CpGs. Capture platforms are more precise and can be customized for profiling specific genomic regions of interest. In addition, capture platforms are the only methods available for methylation profiling of low quality, degraded DNA although library preparation is more labor intensive and efficiency of capture depends on shearing.

## Conclusions

Epigenetic state is a fundamental element of cellular development and regulation. Therefore, accurate, reproducible and cost effective approaches for profiling DNA methylation are important for advancements in biomedical research. While the cost of sequencing continues to decrease, reaching sufficient coverage for reliable measurement of CpG methylation by WGBS is still prohibitive. We conclude from our comparative study that capture-based approaches provide comparable results and cover more precisely their intended targets than ERRBS, which is a restriction enzyme based approach. They also provide the added flexibility of designing custom capture for surveying regions of interest. On the other hand, digestion based protocols are currently more cost effective and may be the only option for clinical samples where the amount of input DNA is limited. In the absence of any prior knowledge about the genomic regions of interest for a particular study, the choice of methylation profiling platform should be guided by cost and amount and quality of input DNA.

## List of abbreviations

5mC: 5-methylcytosine
BS-seq: bisulfite sequencing
CpGiant: Roche NimbleGen SeqCap Epi CpGiant
CpGi: CpG island
ERRBS: enhanced reduced representation bisulfite sequencing
GEO: gene expression omnibus
MAD: median absolute deviation
PCR: polymerase chain reaction
SSMethylSeq: Agilent SureSelect Methyl-Seq
WGBS: whole-genome bisulfite sequencing

## Declarations

### Ethics approval and consent to participate

Not applicable

### Consent for publication

Not applicable

### Availability of data and material

The data that support the findings of this study are available in NCBI GEO (http://www.ncbi.nlm.nih.gov/geo/) under Accession Number GSE83595. The code for the analysis is available from the code hosting platform GitHub (https://github.com/thk2008/methylseqplatformcomparison)

### Authors’ contributions

DB, AA Conceived and designed the experiments. MPF, XZ, YL performed the experiments. TJK performed the bioinformatic analysis and prepared the figures. TJK, DB, AA, analyzed the data. TJK, DB drafted the manuscript. TJK, DB, AA, MPF reviewed the manuscript. All authors read and approved the final manuscript.

## Additional file

Additional file1. **Table S1**. Library input details. **Table S2**. Target region properties and CpGs covered. **Table S3**. Sequencing details. **Figure S1**. Number of CpG-units covered. **Figure S2**. Mean and median coverage per CpG-unit. **Figure S3.** Intra- and Inter- platform CpG-unit overlap and methylation levels concordance. **Table S4**. Intra- and Inter- platform details. **Figure S4**. Overlap of exon annotation of CpG-units as UpSet plot. **Figure S5**. Overlap of intron annotation of CpG-units as UpSet plot. **Figure S6**. Overlap of promoters annotation of CpG-units as UpSet plot. **Figure S7**. Overlap of CpG island annotation of CpG-units as UpSet plot. **Figure S8**. Overlap CpG shores annotation of CpG-units as UpSet plot. **Figure S9**. Overlap of unannotated CpG-units as UpSet plot.

